# Deconstructing the Cancer Epigenome using Reduced Representation Bisulfite Sequencing (RRBS)

**DOI:** 10.1101/2025.05.12.653548

**Authors:** Nicole Glendinning, Dongjun Chung, Gary Hardiman

**Author notes:** Correspondence should be addressed to Gary Hardiman;,.

## Abstract

Reduced Representation Bisulfite Sequencing (RRBS) represents a relatively inexpensive methodology for investigating the influence of DNA methylation on the development and progression of diseases including cancer. RRBS is a targeted approach which combines the use of restriction enzyme selection, bisulfite conversion, and high throughput sequencing technologies. The computational processing and analysis of RRBS data therefore requires modified approaches compared to typical DNA sequencing, due to deviations from standard laboratory protocols. This chapter presents a step-by-step pipeline for the processing of raw RRBS reads including quality control, trimming, alignment, methylation extraction, differential methylation analysis, annotation, and downstream analysis. This pipeline is designed to ensure optimal processing for accurate results and to provide downstream analysis tools that can extract biological insights from the data.

## 1. Introduction

Epigenetics governs gene expression through mechanisms that do not alter the underlying DNA sequence. Epigenetic processes include DNA methylation, histone modification, and control by non-coding RNAs. DNA methylation is crucial for regulating various biological events, including gene expression, development, X-chromosome inactivation, imprinting, and aging, as well as disease. Abnormal regulation is strongly linked to conditions such as psychiatric disorders and cancer. In the context of cancer, aberrant DNA methylation profiles can drive tumor onset and progression. Hypermethylation in the promoter regions of tumor suppressor genes can silence these genes, permitting unchecked cell division. Conversely, global hypomethylation can destabilize the genome and activate oncogenes. As these epigenetic changes can emerge early in oncogenesis, researchers increasingly focus on these methylation alterations both as prospective diagnostic biomarkers and as leads for novel therapeutic strategies that could significantly enhance treatment outcomes.

DNA methylation is the addition of a methyl group to the fifth carbon of cytosine nucleotides, forming 5-methylcytosine. In mammals, most methylation takes place at CpG sites, where cytosine is followed by guanine. These CpG sites frequently cluster in regions called CpG islands, which are DNA segments rich in CpG sites and often overlap with promoter regions. Gene expression can be regulated by DNA methylation through the blocking of transcription factor binding, or the recruitment of additional proteins that influence expression. Generally, methylation at promoter regions is associated with gene silencing ***(1)***. Aberrant DNA methylation patterns within promoter regions can therefore contribute to tumorigenesis through the silencing of tumour suppressor genes. Likewise, the loss of DNA methylation at important promoters can result in inappropriate activation of oncogenes ***(2)***.

Bisulfite treatment of DNA in combination with high throughput DNA sequencing technologies can be used to assess DNA methylation patterns within the genome. Bisulfite treatment is a laboratory method that results in the conversion of unmethylation cytosines in DNA to uracil, whilst methylated cytosines remain unchanged. The resulting sequences can be analysed computationally to determine the positions of methylated and unmethylated cytosines within samples. Although advantageous, carrying out bisulfite sequencing on the whole genome can be extremely costly, especially if a study requires numerous replicates or high sequencing depth and coverage is required. Therefore, Reduced Representation Bisulfite Sequencing (RRBS) ***(3)*** was developed as a more targeted, cost-effective method for performing bisulfite sequencing. RRBS combines the use restriction enzymes (usually MspI) with bisulfite sequencing, to select for and analyse regions of the genome which are rich in CpG sites, such as CpG islands, and promoters. Commercial kits or published methods protocols ***(4)*** are available for the laboratory processing of RRBS samples. However, post-sequencing bioinformatic analysis of RRBS data requires tailored strategies that account for both restriction enzyme digestion and bisulfite conversion to ensure accurate results for downstream analysis and data interpretation. This chapter will focus on the computational methods involved in preprocessing, differential methylation analysis, and further downstream analysis of raw RRBS sequencing reads. A rudimentary understanding of the command line environment will be assumed going forward. Although the command line can initially appear intimidating to new users, there are countless free online learning resources available, and a base knowledge easily be attained given a relatively minor time investment. Furthermore, an awareness of standard sequencing file types would be advantageous; therefore, we direct unfamiliar readers to additional bioinformatics protocols for a richer explanation of raw data files ***(5)***.

## 2. Materials

### 2.1 Computing Resources

To secure the computational resources required for next-generation sequencing data analysis, bioinformatics researchers commonly utilize High Performance Computing (HPC) clusters. Many universities have HPC services available for research staff and students to avail of, doing so can substantially reduce the run times of RRBS data pre-processing steps. Although many bioinformatics tools will function on Windows operating systems, they generally obtain better support for Linux and macOS. If HPC resources are not available; RRBS analysis can run on a personal computer, as most of the tools that will be discussed can also run on macOS, and Windows systems (for Windows operating systems this may require using a Linux environment through software such as Windows Subsystem for Linux https://learn.microsoft.com/en-us/windows/wsl/about). However, users would need to ensure the personal computer has sufficient power, as the main alignment tool discussed requires 5 CPU cores and 16GB ram as a minimum.

### 2.2 Environment Management

As bioinformatic analysis often involves installing numerous different tools, packages, and dependencies; although not essential, it can be extremely beneficial to use system package managers such as Conda, which allows software to be isolated into separate environments. For example, if a system has Python v3.11.0 installed and one of the tools requires Python v3.9.0 for compatibility, it is unnecessary to remove the existing installation to downgrade. Instead, separate environments can be created with different Python versions installed for distinct tasks. This approach prevents conflicts between programs and allows for seamless switching between environments based on the analysis being performed. Conda is available for Linux, macOS, and Windows (https://anaconda.org/anaconda/conda), it is well supported and adopted, with many free online learning resources.

### 2.3 Software

Bismark (https://www.bioinformatics.babraham.ac.uk/projects/bismark/), written in Perl for Linux and macOS. Based on Bowtie2 by default (https://bowtie-bio.sourceforge.net/bowtie2/index.shtml).

BS-Seeker2 (https://github.com/BSSeeker/BSseeker2), Linux and macOS.

bwa-meth (https://github.com/brentp/bwa-meth), Linux and macOS.

Ensembl (https://www.ensembl.org/),Website.

FastQC (https://www.bioinformatics.babraham.ac.uk/projects/fastqc/), Linux, macOS and Windows. Java based.

GeneCards (https://www.genecards.org/), Website.

ggplot2 (https://ggplot2.tidyverse.org/), Linux, macOS and Windows. R package.

g:Profiler (https://biit.cs.ut.ee/gprofiler/gost), Website.

Homer (http://homer.ucsd.edu/homer/index.html), Linux and macOS. Pearl and c++ based.

matplotlib (https://matplotlib.org/), Linux, macOS and Windows. Python Package.

methylKit (https://www.bioconductor.org/packages/release/bioc/html/methylKit.html), Linux, macOS and Windows. R Bioconductor package.

MultiQC (https://seqera.io/multiqc/), Linux, macOS and Windows. Python based.

Python (https://www.python.org/), Linux, macOS and Windows.

R (https://www.r-project.org/), Linux, macOS and Windows.

rnBeads (https://www.bioconductor.org/packages/release/bioc/html/RnBeads.html), Linux, macOS and Windows. R Bioconductor package.

seaborn (https://seaborn.pydata.org/), Linux, macOS and Windows. Python package.

TopFunn (https://toppgene.cchmc.org/), Website.

Trim Galore (https://www.bioinformatics.babraham.ac.uk/projects/trim_galore/), Linux and macOS. Pearl wrapper for Cutadapt (https://cutadapt.readthedocs.io/en/stable/) and FastQC.

UCSC Genome Browser (https://genome.ucsc.edu/cgi-bin/hgGateway), Website.

## 3. Methods

The RRBS methods detailed in this chapter can be separated into three main sections: preprocessing, differential methylation analysis, and downstream analysis (Figure 1). Preprocessing consists of completing quality checks on the raw data, trimming low-quality information, alignment of trimmed reads to a reference genome, and methylation extraction. Differential methylation analysis involves the identification of differential methylation between test and control samples. Lastly, downstream analysis incorporates further exploration of differential methylation patterns within a more biologically relevant context by investigating pathways that may be disrupted, and through visualisation of results. It is important to note that for all RRBS analysis, the documentation associated with laboratory processing should be consulted for computational pre-processing recommendations, as some manufacturers can require the use of additional bespoke parameters.

**Figure 1:**
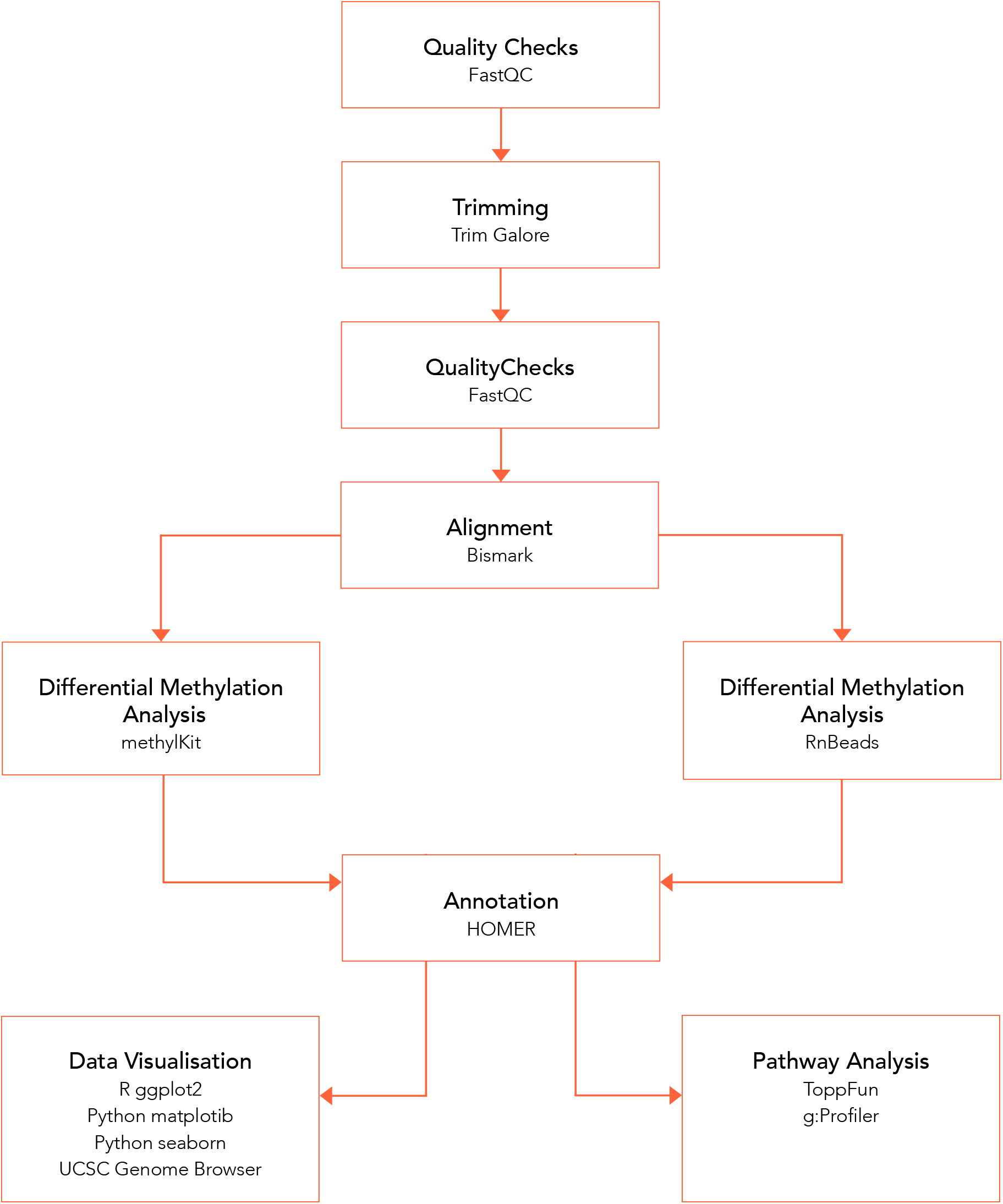
Flow Chart summarising the steps involved in RRBS computational analysis from raw FASTQ files.

In the case of RRBS, two main components require consideration of experimental designs: (i) detection of methylated cytosines/regions and (ii) detection of DMCs/DMRs. In the case of detection itself (factor (i)), the total sequencing depth is the main driver that determines the detection, i.e., we need to determine the minimal total sequencing depth that will guarantee the detection of methylated cytosines/regions. In the case of differential analysis (factor (ii)), both the sample size (number of animals/subjects) and the total sequencing depth matter but in general determining the minimum required sample size is of main interest because it affects the experimental cost more directly. For the first type of power analysis, users can use tools like the Shiny app ‘covcalc’ (https://stephenturner.shinyapps.io/covcalc/). The power analysis for the second type has been investigated more and users can use tools like the Shiny app ‘bisulfitesim’***(6)***; http://www.tung-lab.org/protocols-and-software.html) or the R package ‘MethylSeqDesign’ ***(7)*** (https://github.com/liupeng2117/MethylSeqDesign) for such type of power analysis.

### 3.1 Preprocessing

#### 3.1.1 Quality Checks (QC)

Before performing any downstream analysis on high throughput sequencing data, it is essential to verify that the raw FASTQ files meet high quality standards for any sequencing method. This important step ensures there are no data concerns stemming from technical issues during laboratory processing or sequencing that may need to be considered moving forward. FastQC ***(8)*** is an extensively used program for performing quality checks (QC) of raw sequencing data. The “fastqc” function of FastQC can be used in conjunction with raw FASTQ files to create HTML reports detailing how a single sample performs in relation to numerous metrics such as sequence quality, adaptor contamination, or overrepresented sequences. However, as this can often involve quality checking large numbers of individual FastQC reports, MultiQC ***(9)*** can be used via the “multiqc” command to aggregate numerous FastQC reports into one convenient summary report based on the same metrics.

It is important to note, that as RRBS represents a tailored methodology, FastQC and MultiQC reports may look different to standard exemplar reports, and data might not meet the expectations of the standard QC metrics used for RNAseq and DNAseq. Even in otherwise good quality RRBS QC reports, these tools can often highlight concerns with metrics such as: per base sequence content, per sequence GC content, sequence duplication and overrepresented sequences (Figure 2). Flagged issues with per-base sequence content and per-sequence GC content may simply reflect the choice of restriction enzyme used in RRBS. The use of restriction enzymes such as MspI can result in almost all reads starting with the same base composition ***(10)***; Similarly, because the targeted CpG sites often lie within regions of high GC content, it follows that the GC content of the resulting reads may differ from what is typically observed in standard sequencing. FastQC and MultiQC warnings associated with sequence duplication and overrepresented sequences can similarly be linked to RRBS representing a genome complexity reduction method. As RRBS prepared libraries target specific genomic regions, there will be a smaller pool of potential reads for sequencing, leading to the same regions being covered more frequently than what would be expected through untargeted approaches.

**Figure 2:**
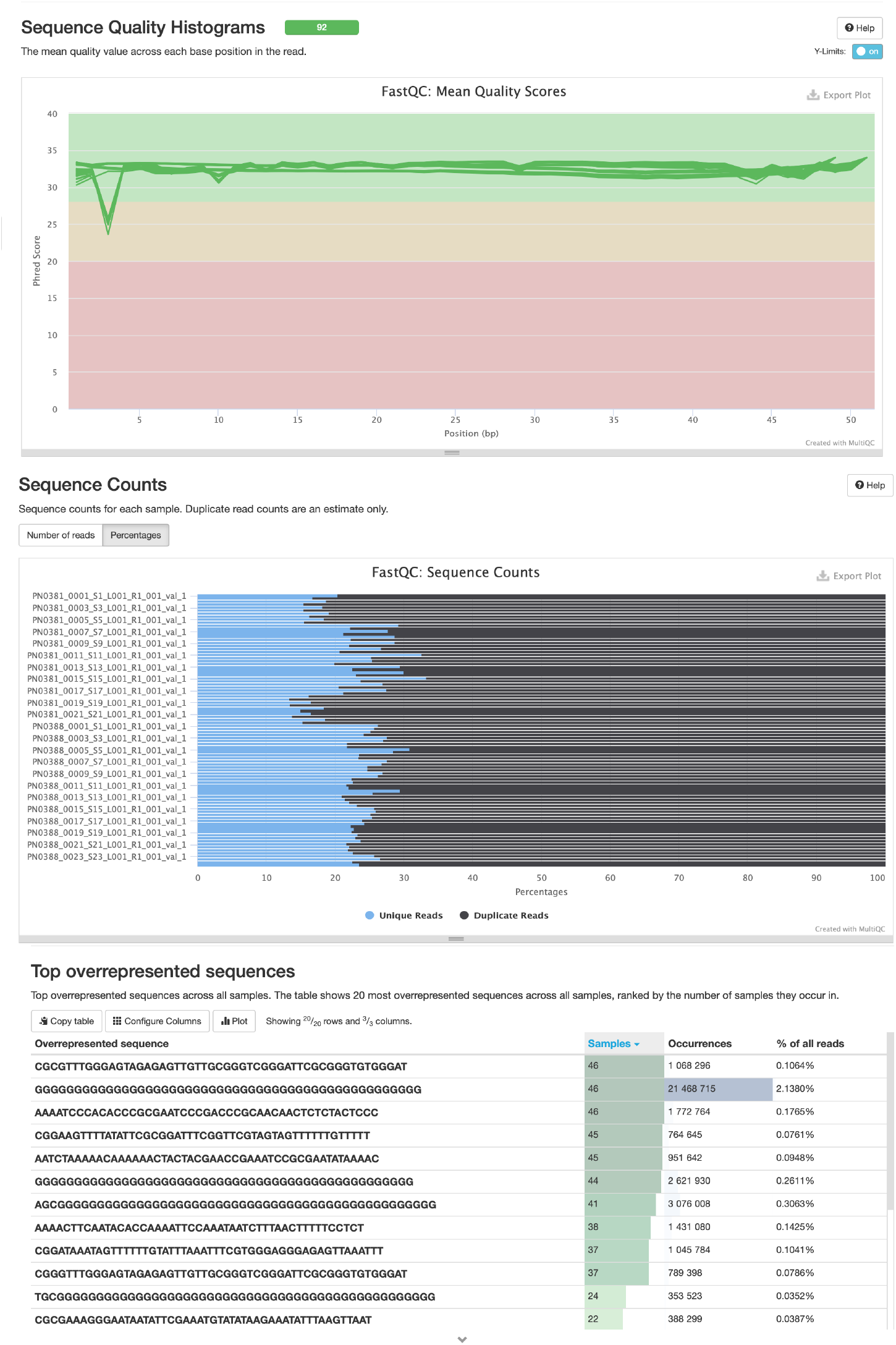
Example RRBS MultiQC report. Highlighting good sequence quality, but high levels of sequence duplication and overrepresented sequences.

Overall, it is important for users to check all metrics to ensure they are satisfied with the data prior to further analysis. Completing a full inspection and evaluation of QC reports is essential to ensure satisfaction with overall sequence quality, and to ensure that any flagged issues are a consequence of the sequencing methodology used and not indicative of more serious issues with prior processing.

#### 3.1.2 Trimming

For all sequencing technologies, following quality checks, concerns with the data such as low-quality sequences or adapter contamination can be improved by carrying out trimming. In addition to this, the Mspl restriction enzyme digestion as part of RRBS can lead to data being prone to additional biases. After Mspl digestion, an end repair reaction results in additional cytosines being introduced, which can either be methylated or unmethylated depending on laboratory processing ***(10)***. To ensure that these extra cytosines do not interfere with methylation calling, additional trimming can be carried out using Trim Galore ***(11)***. Trim Galore is a wrapper for Cutadapt ***(12)*** that allows for standard adapter and quality trimming, as well as the removal of the artificially introduced cytosines which are a more specific feature of RRBS.

The “trim_galore” function will perform automatic adapter trimming by auto-detecting commonly used manufacturer adapter sequences. Quality trimming is optional and can be executed by specifying ‘-q’ along with the desired Phred score to trim bases on. The ‘--rrbs’ parameter should be used with RRBS data to trim experimentally introduced cytosines (*see* **Note 4.1**). Trim Galore produces trimmed FASTQ files which will again need to be quality checked using FastQC/MultiQC before progressing to the next step of analysis.

#### 3.1.3 Alignment to a Reference Genome and Methylation Extraction

Once sequences have been trimmed and QC checked, reads can then be mapped to known positions of a reference genome through alignment. Due to the conversion of unmethylated cytosines during bisulfite sequencing, standard aligners are not appropriate for RRBS analysis. Bismark ***(13)*** is an aligner based on Bowtie2 by default ***(14)*** (*see* **Note 4.2**), which has been specifically designed to accommodate the alignment of bisulfite converted data. In comparison studies with other tools, Bismark performs well both in terms of accuracy and recall ***(15)***. It has been widely adopted for RRBS and whole genome bisulfite sequencing analysis, as well as being well maintained and updated by its active creators.

To perform an alignment, a copy of the reference genome for the species of interest should be downloaded from an online resource such as UCSC, NCBI or Ensembl. The first step in Bismark alignment is creating a bisulfite converted reference genome using the ‘bismark_genome_preparation’ command. Supplying the path to the downloaded reference genome to this command will create two additional bisulfite converted versions of the genome, a C>T converted reference, and a G>A converted reference.

Next, the ‘bismark’ command can be used to perform alignment to the reference genome. This command requires the path to the folder containing the reference genomes, along with the trimmed and QC checked FASTQ files for alignment. The resulting output will be in BAM (Binary Alignment Map) file format (Figure 3), (*see* **Note 4.3 and Note 4.4**).

**Figure 3:**
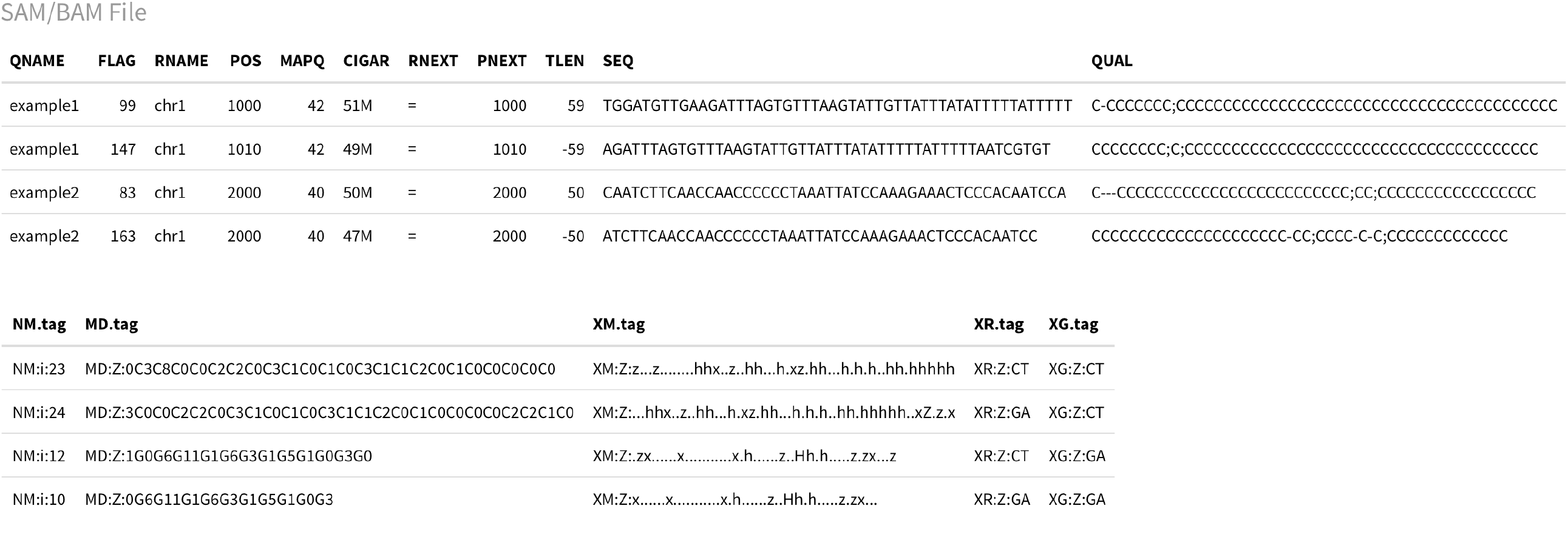
Simplified example of Bismark alignment file outputs. The BAM (Binary Alignment Map) files is the compressed form of SAM (Sequence Alignment Map) files. Further information on the fields included in a Bismark BAM/SAM file are detailed within the Bismark online documentation available at: https://felixkrueger.github.io/Bismark/bismark/alignment/. [Accessed 11/02/2025].

Lastly, Bismark’s ‘bismark_methylation_extractor’ command can be used to extract methylation calls from the aligned BAM files. Providing the ‘--bedgraph’ parameter to this command will save outputs as bedGraph and Bismark coverage files (.cov files) (Figure 4). Both the bedGraph and Bismark coverage files detail the genomic position of each cytosine site analysed as well as a percentage methylation score associated, with the Bismark coverage files containing additional coverage information. Although the methylation extraction step is not strictly necessary, many downstream tools have infrastructure in place to easily read in and use Bismark coverage files without additional cleaning or data manipulation steps.

**Figure 4:**
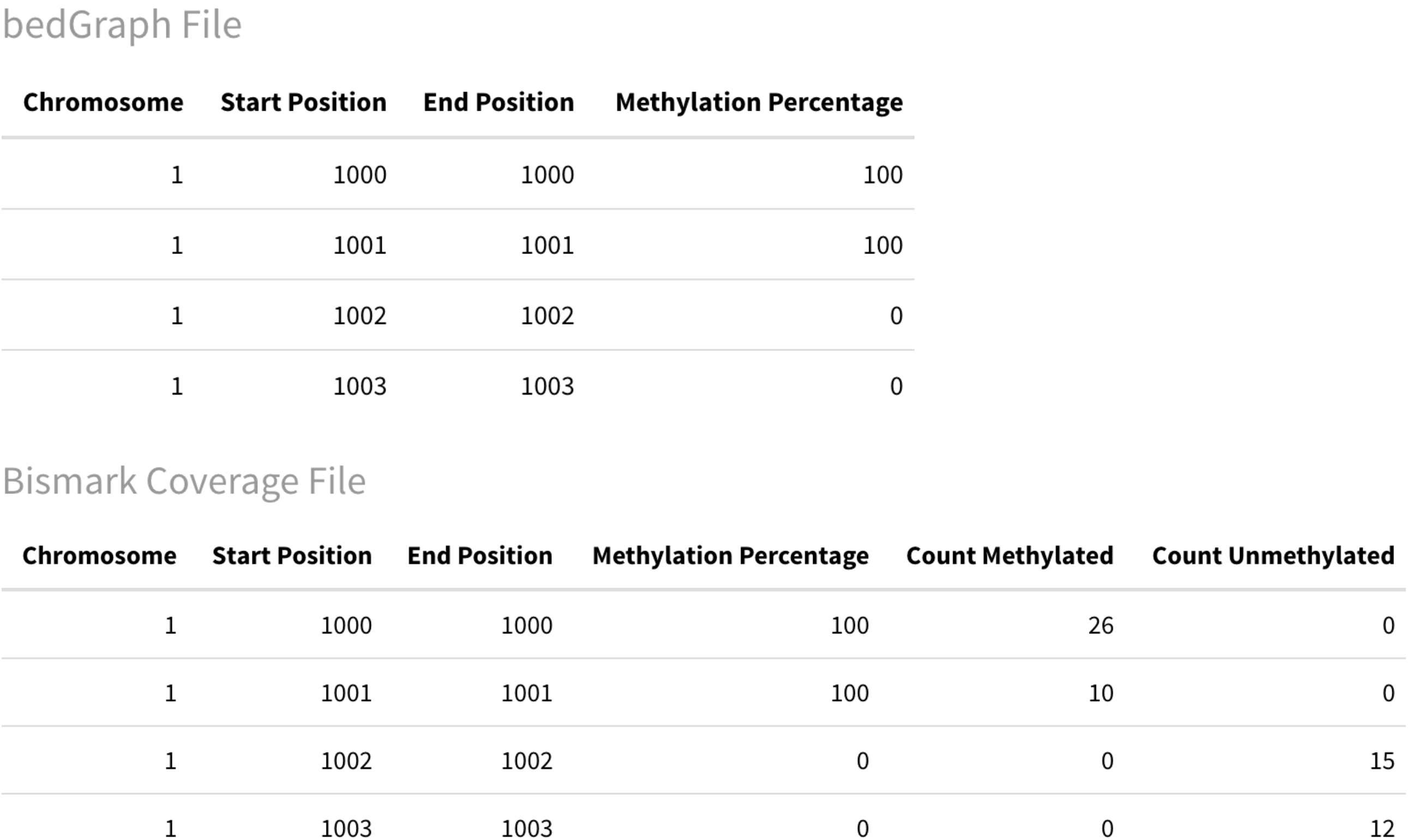
Simplified example of bedGraph and Bismark Coverage format produced by methylation extraction. Bismark Coverage files provide additional coverage information through the ‘Count Methylated’ and ‘Count Unmethylated’ columns.

MultiQC can again be utilised in combination with Bismark outputs to produce summary alignment and extraction QC reports. These QC reports can be assessed to examine the percentage of reads that Bismark has aligned, and overall methylation levels for samples being processed. This again gives users the opportunity to assess if these metrics are in line with their expectations given the data and can highlight potential problems with processing.

### 3.2 Differential Methylation Analysis

Differential methylation analysis involves the identification of differentially methylated cytosines (DMCs), or differentially methylated regions (DMRs) between test samples, and control samples. Two widely used R Bioconductor packages that facilitate differential methylation analysis are methylKit ***(16)***, and RnBeads ***(17)***.

#### 3.2.1 methylKit

MethylKit has extensive and clear documentation available for new users, it is extensively used and has shown good performance under a range of different circumstances in comparison studies ***(18), (19)***.

Files are read into methylKit via the “methRead” function; lists of Bismark coverage files can be supplied directly to this function by indicating “pipeline=‘bismarkCoverage”. Test and control samples must be specified by providing a vector of 1’s and 0’s to the “treatment” parameter, the order of this must correspond to the order in which the sample files are provided. By default, methylKit requires a minimum of 10X coverage at each cytosine site and sites not meeting this threshold will not be included in the analysis; this can be altered using the “minicov” parameter (*see* **Note 4.5**). methylKit will store this information as a “methlRawList” object, which contains methylation information on cytosines meeting coverage thresholds per sample indicated.

Next, the “unite” function is employed to merge the numerous individual samples within the “methlRawList” from the previous step into one larger R “methylBase” object containing cytosine sites which are covered in all samples.

Lastly, DMCs are identified by suppling the united “methylBase” object to the “calculateDiffMeth” function. This will automatically calculate p-values and adjusted p-values, as well as the percentage methylation difference between test and control samples. methylKit will either use Fisher’s exact test, or logistic regression to calculate differential methylation, depending on the number of samples included in the analysis. Adjusted p-values will be calculated using a SLIM method by default; this can be updated to use False Discovery Rate (FDR) by specifying “adjust=“BH”“. The results can be filtered based on user-defined significance thresholds and saved as a CSV file for further downstream analysis. Although methylKit does not produce extensive figures like RnBeads, results are clean and comprehensible as shown in Figure 5, as well as being well formatted for use with additional downstream tools.

**Figure 5:**
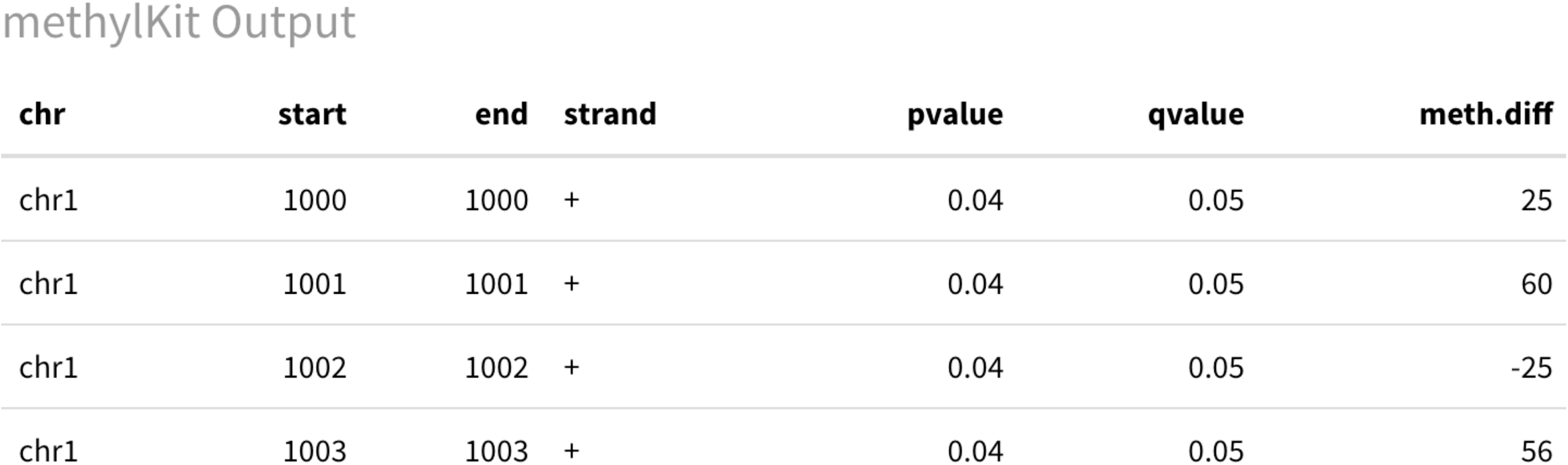
Simplified example of methylKit DMC results format.

MethylKit can identify DMRs via the same functions as DMCs, with slight modifications. As DMRs consider much larger regions of the genome than individual DMCs, when reading in files with the “methRead” function, users may wish to reduce the minimum coverage per cytosine requirement from the default value of 10X to e.g. 3X through the “minicov” parameter. After files have been read into methylKit and stored a “methlRawList” object, the “tileMethylCounts” function can be applied to split the genome into regions of default 1000bp and summarise the methylation status of the cytosines covered within the whole region per sample. After the tiling function is applied, samples can be united and differential methylation calculated as detailed above.

#### 3.2.2 RnBeads

RnBeads supports differential methylation analysis from both microarray-based methods, and bisulfite sequencing methods including RRBS. As part of RnBeads analysis, many high-quality figures, charts and tables are produced and can be collated within a convenient HTML format for easy navigation and review (see Figure 6 for examples). Before analysis can begin, a spreadsheet-like sample annotation file in the form of a text file must be collated and supplied to RnBeads. This annotation file should detail important contextually relevant information such as sample identifiers, full file names of the appropriate sample files, a description of samples (i.e. if sample is test or control), and covariates to be included in analysis (see Figure 7 for a simplified example).

**Figure 6:**
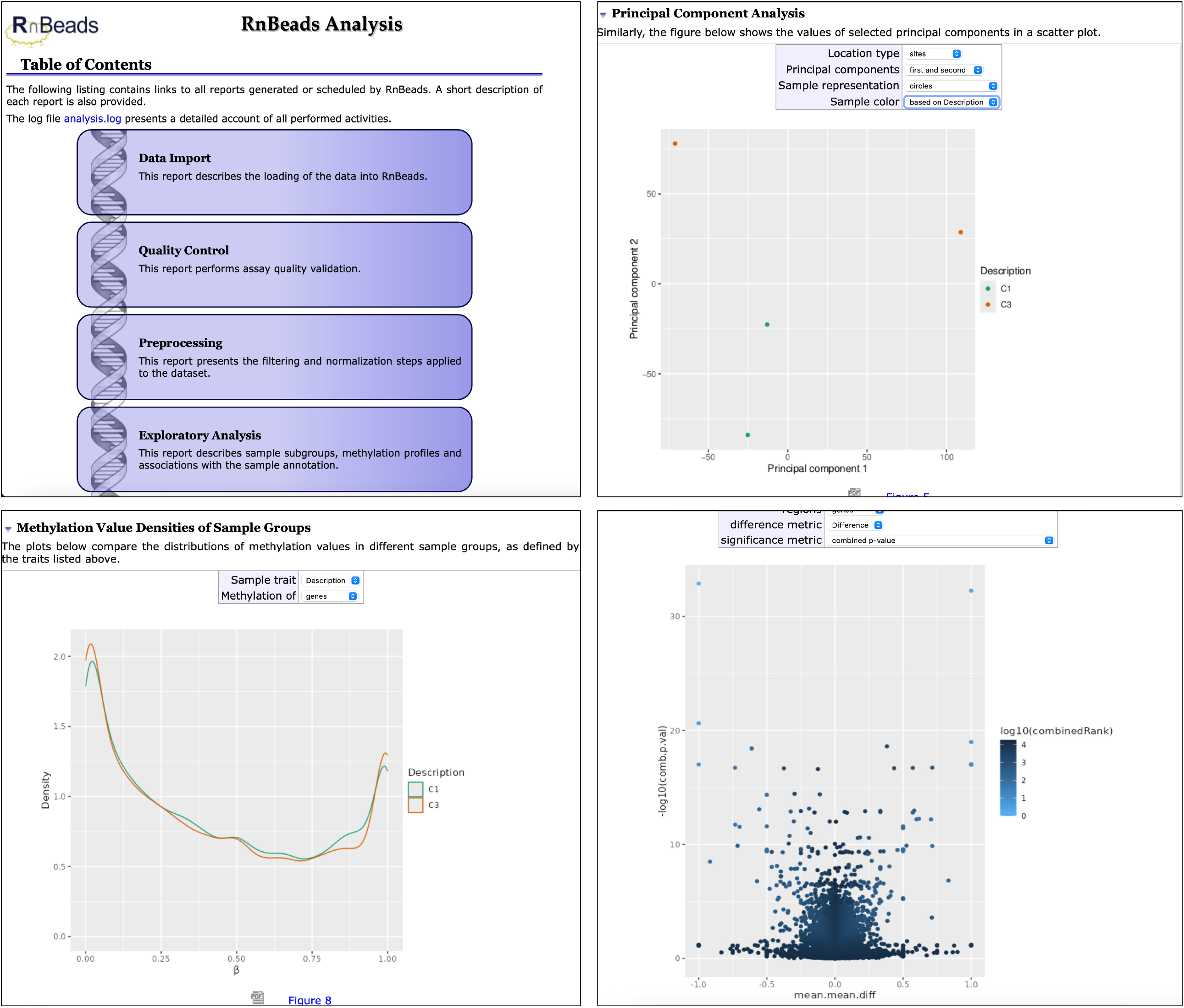
Screenshots of an example RnBeads HTML report for RRBS data. Many of the plots created as part of the standard analysis have modifiable parameters that allows users to complete simple adjustments such as changing the genomic region of interest or adding sample identifiers.

**Figure 7:**
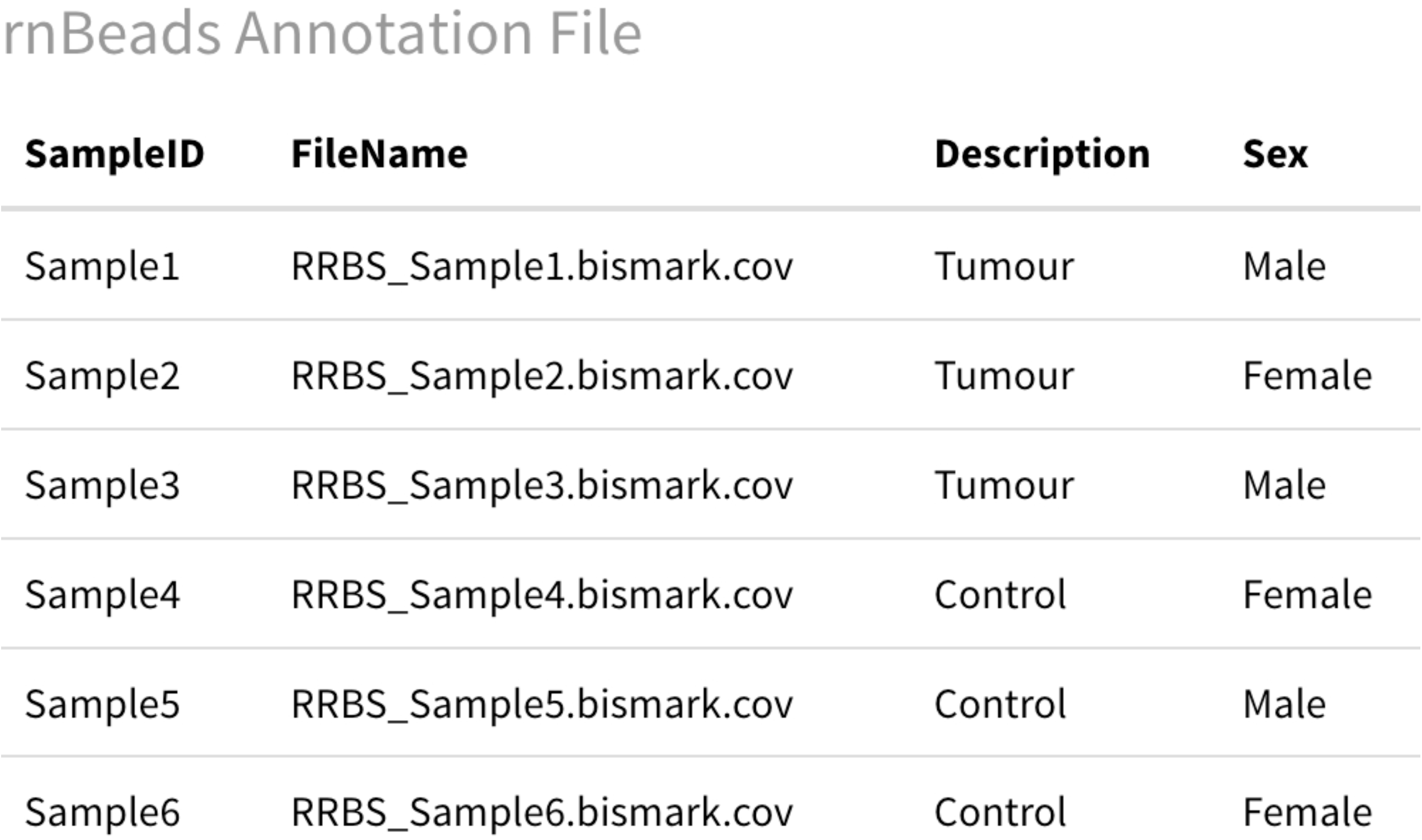
RnBeads exemplar annotation file containing contextual information regarding samples for analysis.

The simplest way to run RnBeads is via what is the authors refer to as a vanilla analysis, where parameters are declared, and the entire RnBeads pipeline is run on the data in one pass. This will output all data and figures available as part of RnBeads.

The first step of this vanilla analysis involves declaring specific parameters via the “rnb.options” function. There are an extremely large number of parameters associated with this function and for the purposes of this chapter, many will be left as default, with essential options that must be updated for basic RRBS analysis detailed (*see* **Note 4.6**). Similarly to methylKit, Bismark coverage files can be read directly into RnBeads, this file type simply needs to be indicated during parameter setting by specifying “import.bed.style= “bismarkCov”“. During parameter setting, additional information concerning the annotation spreadsheet will need to be supplied as RnBeads will refer to this file during analysis. The sample ID column name from the annotation file can be specified using “identifiers.column”. The column that you want to compare (e.g. indicating if sample if tumour or control) should be indicated via “differential.comparison.columns”. Additionally, as part of RnBeads analysis, preliminary annotation of results with gene and promoter information is supported. This feature relies on RnBeads annotation packages which are available within Bioconductor for supported genome assemblies (currently hg19, hg38, mm9, mm10 and rn5), and can be installed via R Bioconductor prior to analysis (*see* **Note 4.7**). The “rnb.get.assemblies()” function can be used in order to determine which assemblies are present within the system, and the genome assembly of interest can then be indicated during option configuration using the “assembly” flag.

Following configuration, the analysis itself is run via the “rnb.run.analysis” function. For all bisulfite sequencing analyses “data.type=“bs.bed.dir”” should be specified within this function. Additionally, indicate via “dir.reports” the full file path to the folder where the output should be saved, as well as well as the path to the sample annotation file using “sample.sheet”. The directory containing samples for processing can be supplied to the “data.dir”, parameter.

RnBeads results will be saved in numerous different CSV files for the application of additional filtering based on user defined thresholds. However, it is important to note, that as RnBeads results are broken into multiple spreadsheets which each contain many columns; understanding, filtering and cleaning this data into a format appropriate for downstream applications may require a greater time investment compared to methylKit results.

### 3.3 Annotation

To obtain additional functional information regarding the areas of the genome DMCs and DMRs reside in, additional annotation tools such as HOMER ***(20)*** can be employed. HOMER includes the “annotatePeaks.pl” program which can perform functional annotation of genomic sites or regions highlighted by methylKit or RnBeads. The data returned by HOMER includes gene identifiers, if the site/region resides in a intron, exon or intergenic region, as well as information regarding the nearest transcription start site. The data for annotation can be formatted as a variation of a BED file which is a tab delimited text file containing each DMC/DMRs genomic location, as well as a unique identifier which authors refer to as a Peak ID (Figure 8) (*see* **Note 4.8**). HOMER contains built in support for numerous genome versions including human hg38, mouse mm10, and rat rn6; the genome will need to be specified when running “annotatePeaks.pl” (*see* **Note 4.9**).

**Figure 8:**
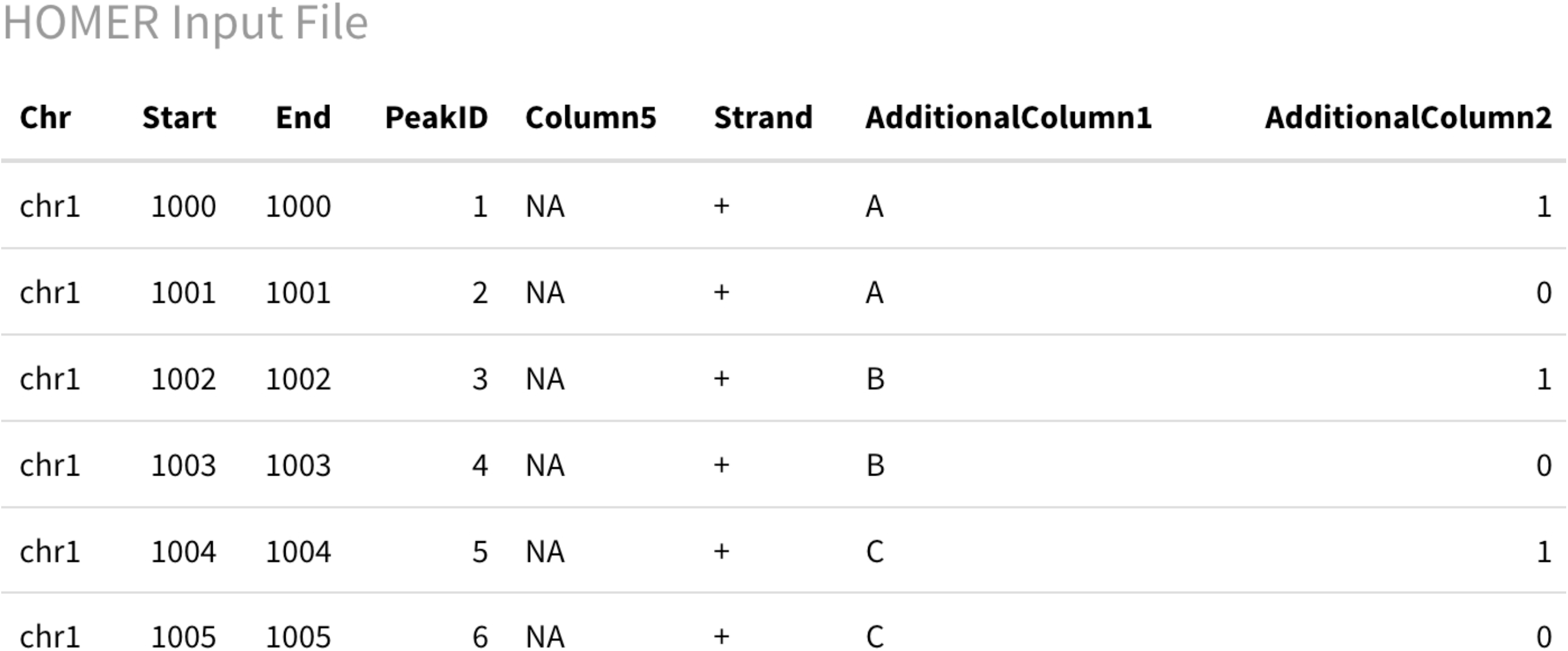
Example HOMER BED Input file. HOMER requires a minimum of 6 columns: chromosome, start position, end position, a unique ‘PeakID’ identifier, a placeholder Column5, and strand information. Spreadsheets can have additional columns after these essential 6.

### 3.4 Pathway Enrichment Analysis

From annotated DMC and DMR results, additional information on associated genes can be compiled from the web-based databases such as GeneCards ***(21)*** or Ensembl ***(22)***. These databases can provide information such as potential function, protein structure, associated disorders, as well as pointing viewers towards PubMed IDs of papers relevant information has been pulled from.

Although it can be extremely beneficial to analyse DMC and DMR results individually, depending on the filtering thresholds defined this could potentially lead to hundreds of independent genes to evaluate. Therefore, it may be more feasible to complete full searches for a small subsection of top hits returned, and to focus predominantly on pathway enrichment analysis. Pathway enrichment analysis can assist with condensing large lists of DMC and DMR related genes, into reduced lists of overrepresented pathways. This can help to explain the biological processes or functions that may be impacted in test samples compared to controls.

Free web-based tools for performing pathway enrichment analysis include g:Profiler ***(23)***, and ToppFun ***(24)***. Both tools are intuitive, easy to use, and receive regular updates ***(25)***; an essential feature which ensures results and interpretations are based on up-to-date material within the field ***(26)***. For ToppFun and g:Profiler, lists of genes can be pasted directly into a query box, ToppFunn will assume that human gene symbols have been entered, whereas g:Profiler allows users to select the organism that is under study. Both tools will compare the genes entered with a list of background genes to identify pathways that are significantly enriched in the gene list compared to what would be expected given the background set (Figures 9 and 10).

**Figure 9:**
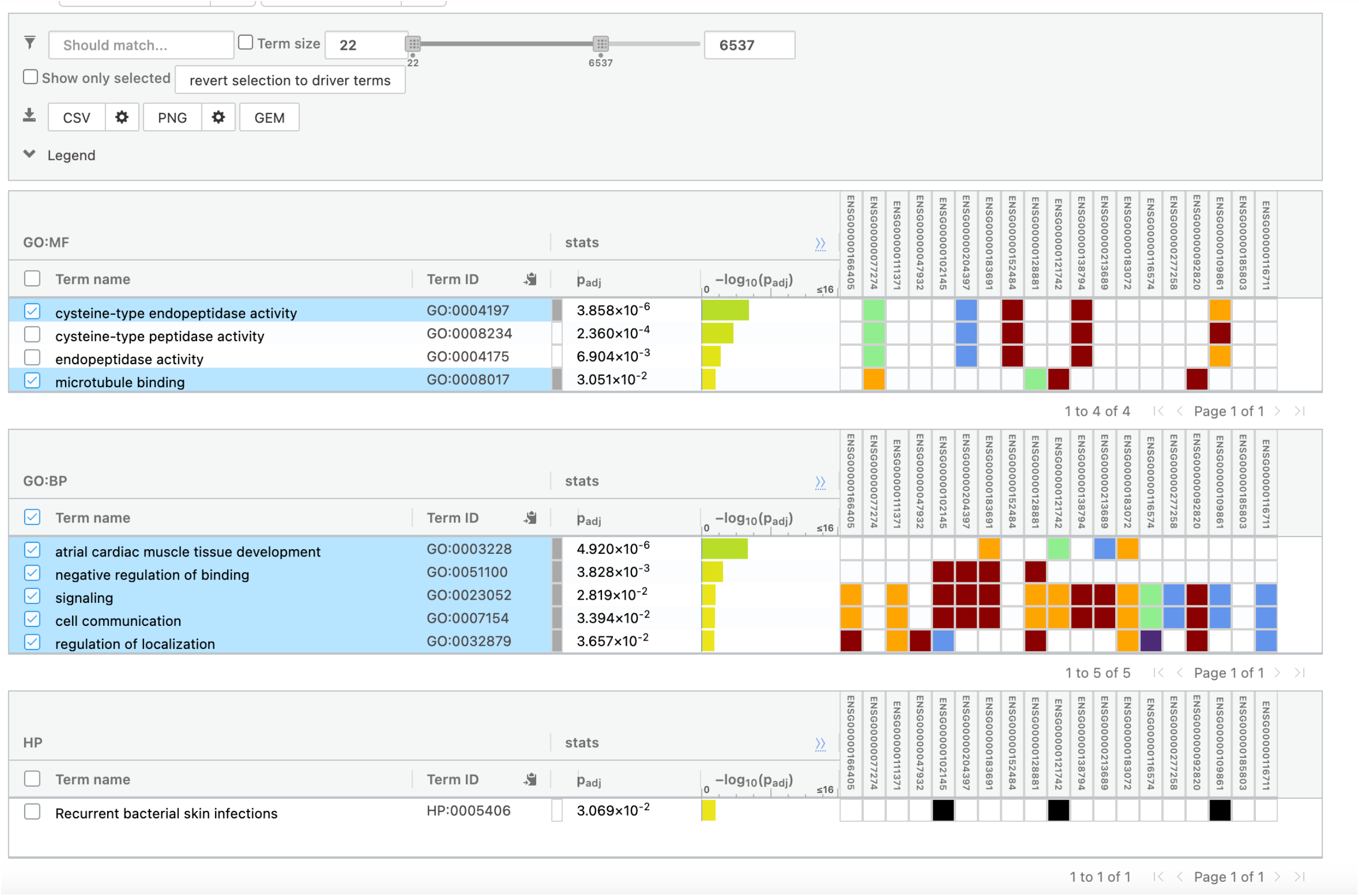
Example g:Profiler results obtained through the ‘Random Example’ option on the g:Profiler homepage. [Accessed 30/01/2025].

**Figure 10:**
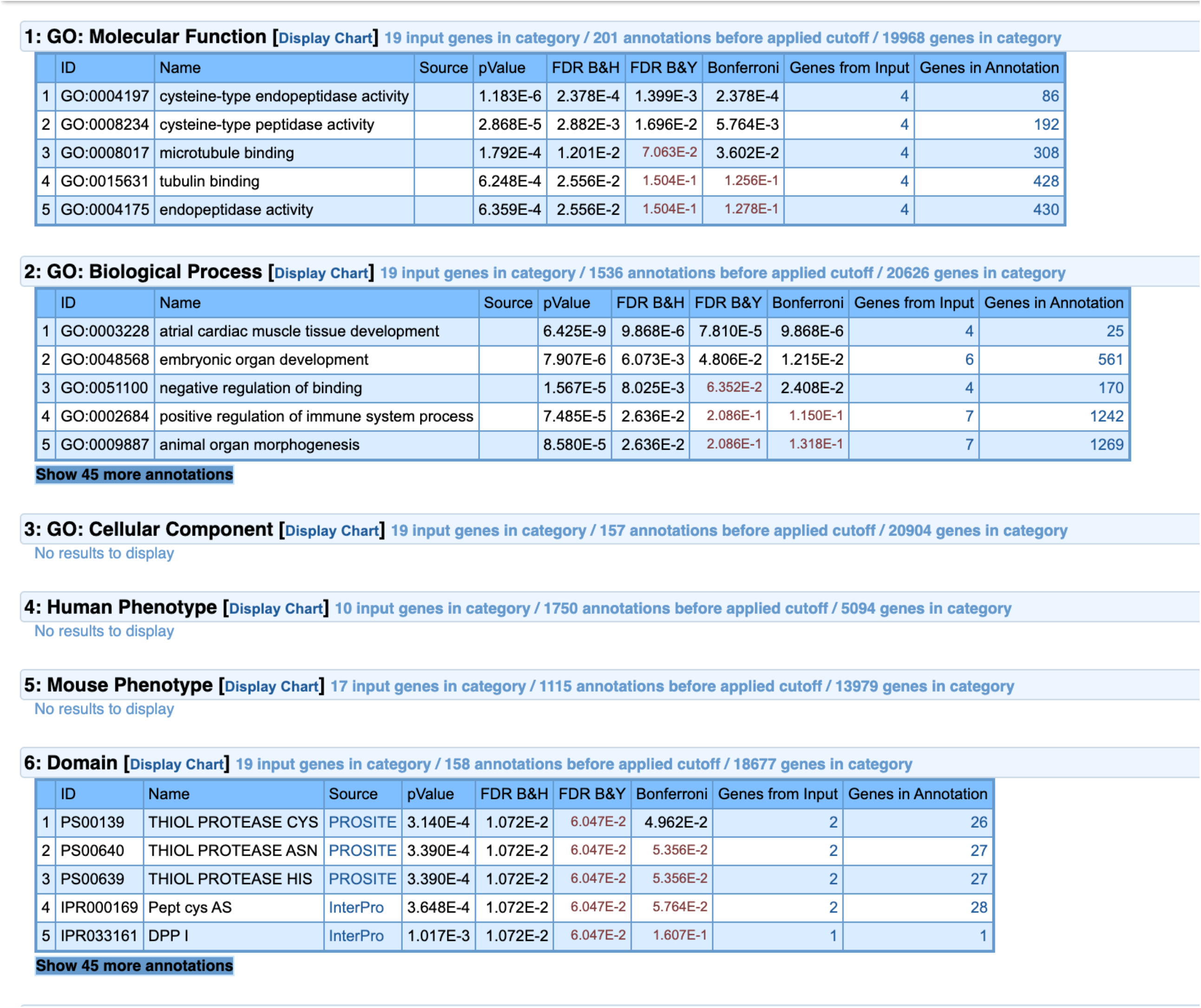
Example ToppFun results given the same g:Profiler ‘Random example’ gene IDs from Figure 9.

### 3.5 Data Visualisation

As raw differential methylation analysis returns potentially thousands of rows in large tables, it can be useful to summarise and communicate main patterns and trends of the data graphically. Even if using the RnBeads pipeline, which produces many figures automatically, it is advantageous to be able to produce more customised figures from filtered results manually.

If there are specific regions of interest, or areas which have a relatively high density of DMCs/DMRs, it may be useful to visualise such regions within genomic space through genome browsers. UCSC Genome Browser ***(27)*** is a web-based tool that allows users to display various types of annotation data such as genes, CpG islands, and repeat regions in relation to genomic position. Within UCSC Genome Browser, users can add their own custom annotations known as tracks, through the “add custom tracks” button in the browser. UCSC will accept custom annotations in multiple different formats; the simplest way to provide DMC or DMR information is through bedGraph files, as these can be produced as an output of methylKit analysis. methylKit can produce bedGraph files by supplying the DMC or DMR outputs to the “bedgraph” function. This function also requires a “col.name” parameter to be set (setting this as “meth.diff” will allow display of methylation percentages), and a “file.name” to be indicated to write the file. When bedGraph files are supplied to the genome browser, DMCs/DMRs positions can be identified visualised in relation to the genomic position, along with the methylation difference (Figure 11) and a range of user defined annotations. UCSC Genome Browser supports human reference genomes, as well as many model organisms.

**Figure 11:**
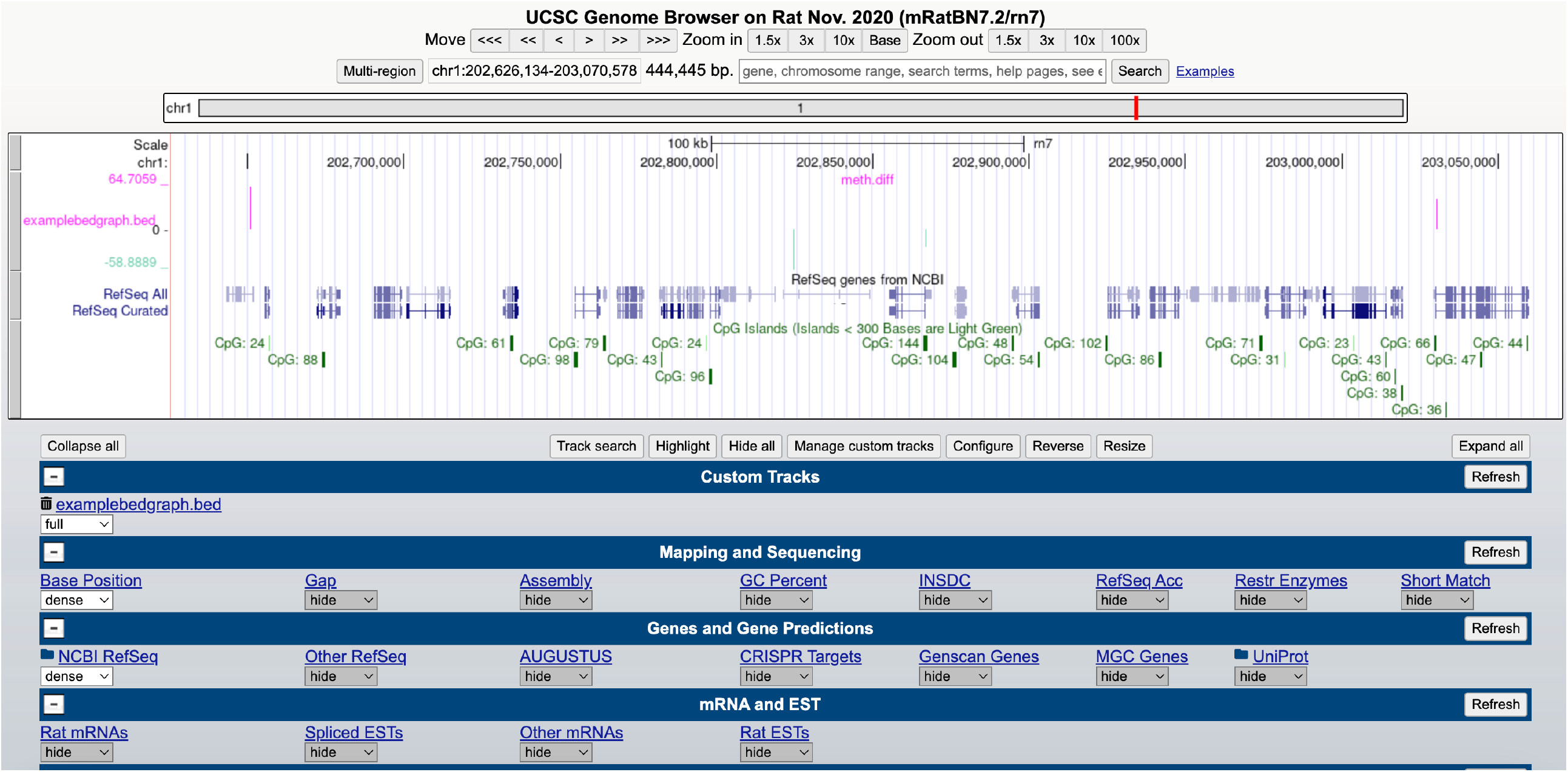
Example UCSC Genome Browser screenshot, showing methylation difference (positive methylation difference in pink, negative methylation different in green) from methylKit, RefSeq gene information, and CpG islands in relation to genomic position.

Additionally, widely adopted packages for data visualisation are available in both R (ggplot2 ***(28)***) and Python (matplotlib ***(29)***, and seaborn ***(30)***). They can be used to make a large variety of engaging plots, and their online documentations contains in depth tutorial with exemplar code, as well as recommended learning resources.

By means of illustration, pie charts can be generated to efficiently summarise the genomic locations within which DMCs and DMRs occur, or bar charts can be used to efficiently display the most significant disrupted pathways from g:Profiler or ToppFun results (Figure 12).

**Figure 12:**
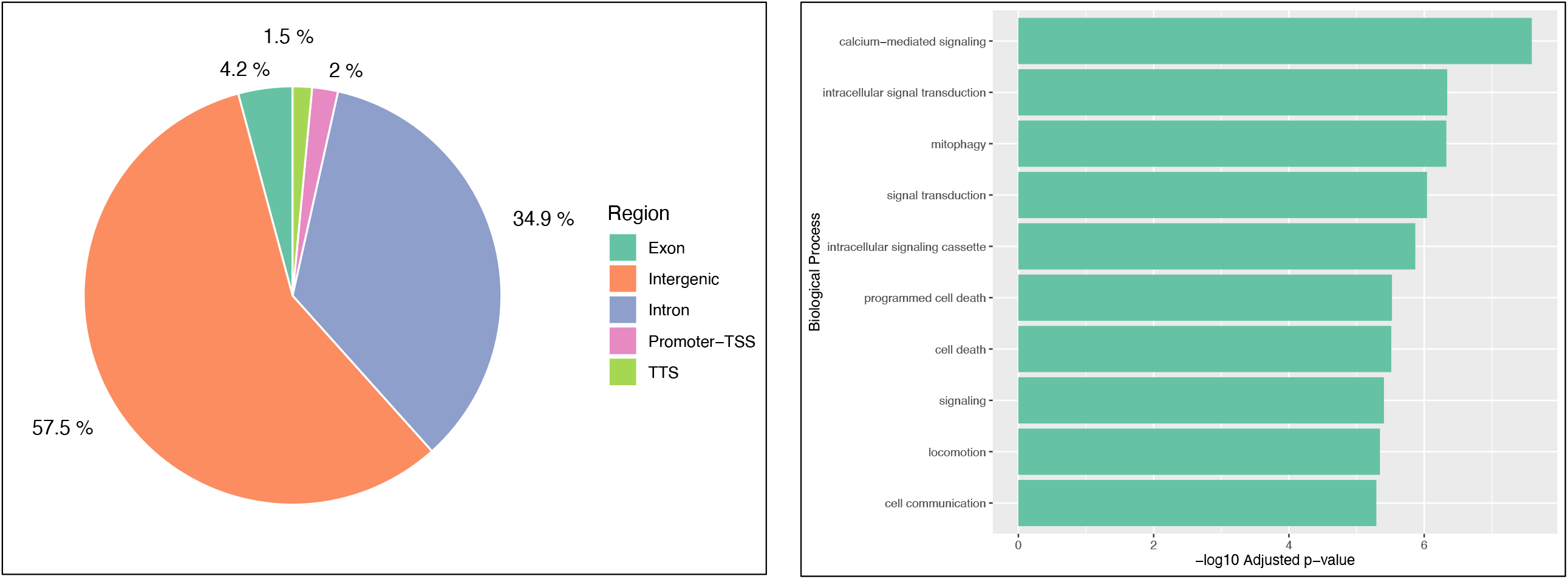
Example data visualisation created using the R ggplot2 package. Example data includes a pie chart summarising locations of DMRs, and a bar chart showing the top enriched biological pathways associated with a gene set.

## 4. Notes

4.1 Check documentation of any commercial kits used to see if reads produced are directional, or non-directional. Non-directional reads have an additional ‘--non-directional’ parameter within Trim Galore.

4.2 Bismark is also compatible with HISAT2 ***(31)*** for alignment instead of Bowtie2. To use HISAT2, the “--hisat2” option needs to be included both during genome preparation, and Bismark alignment.

4.3 Similarly to Trim Galore, if the RRBS reads under analysis are non-directional, an additional ‘--non-directional’ parameter can be supplied to the ‘bismark’ command

4.4 As Bismark does have comparatively longer run times than other tools, it may be worthwhile exploring alternative software if HPC resources are not available; especially if there are a larger number of samples for alignment. BS-Seeker2 ***(32)***, and bwa-meth ***(33)*** have both been recommended by a comparative evaluation of aligners for RRBS data. Both maintain good recall and accuracy, whilst supporting a shorter run time than Bismark ***(15)***. However, additional steps may be necessary to format results appropriately for downstream tools.

4.5 MethylKit provides functions for additional optional descriptive statistics, filtering, normalisation, overdispersion adjustment, and correction for covariates. The steps discussed in this chapter are the essential steps to run differential methylation analysis through methylKit. methylKit documentation (https://bioconductor.org/packages/release/bioc/vignettes/methylKit/inst/doc/methylKit.html) compiles full information on all supplementary steps, and ideally should be consulted prior to running analysis.

4.6. The full list of options for RnBeads is available within the online reference manual (https://www.bioconductor.org/packages/release/bioc/manuals/RnBeads/man/RnBeads.pdf) and should be consulted if deviating from a basic run through the pipeline.

4.7 If analysis involves model organisms, RnBeads may not natively support the most recent version of reference genomes. In this case, unsupported genomes can be used, but it will require the construction of a novel RnBeads annotation package using the “RnBeadsAnnotationCreator” source code available on GitHub https://github.com/epigen/RnBeadsAnnotationCreator/tree/master.

4.8 If using Excel on a macOS system to edit files for Homer Input, text files must be saved as “MS-DOS Formatted Text (.txt)”. Saving the file “Tab-delimited Text (.txt)” within a macOS environment can create issues for the HOMER software.

4.9 If the genome of a model organism used is not natively supported, HOMER can still complete a partial annotation. In this case, the full reference genome FA file along with a GTF annotation file (indicated using the “-gtf” flag) will need to be supplied to the program.

